# STING agonists drive recruitment and intrinsic type I interferon responses in monocytic lineage cells for optimal anti-tumor immunity

**DOI:** 10.1101/2024.12.08.627385

**Authors:** Melanie Girard, Tianning Yu, Nathalia V Batista, Karen K. M. Yeung, Sara Lamorte, Wenting Gao, Miaoxi Liu, Tracy L. McGaha, Tania H. Watts

## Abstract

The cyclic GMP-AMP synthase-stimulator of interferon genes (cGAS-STING) pathway, a sensor of cytosolic DNA, orchestrates the production of pro-inflammatory cytokines, chemokines, and type I interferons (IFN-I), thereby contributing to spontaneous tumor surveillance. Intratumoral delivery of synthetic STING agonists induces IFN-I dependent tumor regression in preclinical cancer models and is being tested clinically. In this study, we investigate the role of monocytic lineage (MCs) cells in response to STING agonist induced IFN-I signaling. We show that CCR2-deficient mice, lacking inflammatory MCs in the periphery, or Lyz2-Cre-IFNAR1^fl/fl^ mice in which IFN-I signaling in monocytes is reduced, exhibit impaired responses to STING agonist therapy of MC38 and/or B16F10 tumors. STING agonist treatment induced CCR5-dependent migration of MCs carrying tumor antigen from the tumor to the lymph nodes. Single-cell RNA sequencing of CD45**^+^** cells from lymph nodes and tumors of mice in which half the hematopoietic cells lack the interferon alpha/beta receptor 1 (IFNAR1) revealed that STING agonist therapy induces intrinsic IFNAR1-dependent acquisition of an inflammatory monocytic cell phenotype distinct from inflammatory cDC and a reduction in macrophages with a pro-tumor TGFβ/angiogenesis transcriptome. IL-18 - IL-18R1 interaction was the top predicted interaction between monocytic lineage cells and CD8**^+^**T cells or NK cells. Blocking IL-18 reduced IFN-γ production by CD8 T cells in LNs and decreased the therapeutic efficacy of STING agonist treatment in CCR2^+/+^ but not in CCR2^-/-^ mice. These findings support a pivotal role for IL-18 producing inflammatory monocytic lineage cells in CD8 T cell control of melanoma following STING agonist treatment.

## Introduction

Monocytic lineage cells (MC), including monocytes, monocyte-derived dendritic cells (DCs) and monocyte-derived macrophages, are highly plastic innate immune cells that adopt distinct functional features depending on the extracellular cues they receive. When MCs enter tumors, they respond to signals from the tumor microenvironment (TME) and differentiate into tumor associated macrophages (TAMs), where they promote tumor progression by suppressing anti-tumor T cell and NK cell responses, and by increasing angiogenesis, tumor cell proliferation and metastasis (1–3). In other contexts, such as during acute viral infections, MCs enter inflamed tissues and respond to proinflammatory cytokines such as type I interferons (IFN-I). These inflammatory monocytic lineage cells (infMCs) help mount effective innate and adaptive immune responses through antigen presentation, co-stimulation, and cytokine production (4–9).

The cyclic GMP-AMP synthase-stimulator of interferon genes (cGAS-STING) pathway elicits an immune response upon the detection of cytosolic DNA (10–13). Upon binding of double-stranded DNA, cGAS catalyses the formation of 2’3’ cyclic GMP-AMP (cGAMP) from cytosolic ATP and GTP molecules (12, 13). cGAMP then binds to the STING adaptor located in the endoplasmic reticulum membrane. This activates NF-κB and IRF3 transcriptional pathways, leading to the production of IFN-Is and other pro-inflammatory cytokines that drive immune responses against viral infection and cancer (10–13).

Sensing of tumor cell-derived DNA via the cGAS-STING pathway in antigen presenting cells is required for spontaneous priming of anti-tumor CD8**^+^**T cells (14). Moreover, DNA sensing through STING and IFN-I production is required for anti-tumor immune responses induced by radiation therapy and chemotherapy (15). These findings prompted the development of small molecules that could activate the STING-IFN-I pathway for cancer immunotherapy. Activation of STING-IFN-I pathway for cancer immunotherapy has shown promising results in pre-clinical mouse models (16–23). However, to date, clinical trials of STING agonists as monotherapy have shown disappointing results (24), highlighting the need to fully understand the mechanisms of protection in the mouse models to facilitate design of optimized combination therapies. ADU-S100 is a synthetic STING agonist that binds to and activates mouse and human STING and leads to downstream IFN-I production (16, 18). Intra-tumoral administration of ADU-S100 induces tumor regression and systemic protection against tumor re-challenge in mice (16, 18). This effect is mediated through CD8**^+^** T cells and/or NK cells depending on tumor-specific features such as MHC class I expression (16, 21). BATF3**^+^** DCs and IFN-I signaling in cDCs are also required for optimal response to treatment, due to their role in cross-presentation and IL-15 trans-presentation (16, 21). There is a well-established role for IFN-I signaling in the response to STING agonist therapy, as IFNAR1 deficiency or IFN-I neutralization significantly impairs tumor control, CD8**^+^** T cell and NK cell responses upon treatment (16, 18, 21).

Depletion of phagocytic cells using clodronate liposomes can reduce the therapeutic efficacy of STING agonist therapy, suggesting that monocytes and/or macrophages are required for the anti-tumor effects of STING agonists (23, 25, 26). Tumor monocytes accumulating in tumors show transcriptional changes following intra-tumoral STING agonist administration (23, 25, 26). Moreover, monocyte depletion impairs STING-dependent radiation therapy (26). However, to what extent these changes are driven by intrinsic IFN-I signaling and whether IFN-I responses in monocytic lineage cells are required for the therapeutic efficacy of synthetic STING agonists has not been directly tested.

To address these questions, we investigated the role of intrinsic IFN-I signaling in MCs during intra-tumoral STING agonist therapy of B16F10 tumors using single cell sequencing and multiparameter flow cytometry. We report that intra-tumoral injection of ADU-S100 drives the accumulation of CD26^lo^CD11b**^+^**Ly6C^hi^CD64**^+^**MAR-1**^+^**infMCs in the lymph nodes (LNs) and tumors of mice. CITE-sequencing of LN and tumor cells from IFNAR1 mixed bone marrow chimeric mice shows that infMCs that accumulate following STING agonist administration have a transcriptional program almost entirely dependent on intrinsic IFN-I signaling and characterized by high expression of IL-18 and genes involved in MHC I-restricted presentation. *Ccr2^-/-^* and *Ifnar1^fl/fl^ Lyz2-Cre* mice show impaired response to STING agonist therapy compared to littermate controls, suggesting that MC and MC intrinsic IFN-I responses are required for optimal response to treatment. Mechanistically, we found that neutralization of IL-18 reduced the therapeutic efficacy of STING agonists in *Ccr2^+/+^* but not *Ccr2^-/-^* mice and decreased CD8 T cell production of IFN-γ, highlighting the importance of this monocyte-derived pro-inflammatory cytokine during STING agonist therapy of cancer. Taken together, these data suggest that recruitment of MCs and IFN-I-induced IL-18 production by infMCs are pivotal for CD8 T cell control of melanoma following STING agonist therapy.

## MATERIALS AND METHODS

### Mice

C57BL/6 and CD45.1 mice were purchased from Charles River Laboratories. *Ifnar1^-/-^*, *Ccr2^-/-^, Ifnar1^fl/fl^* and *Lyz2-Cre* mice were purchased from The Jackson Laboratory. *Ccr2***^+^***^/-^* mice were bred to generate *Ccr2***^+^***^/^***^+^** and *Ccr2^-/-^* littermate controls. *Ifnar1^fl/fl^* and *Lyz2-Cre* mice were crossed to generate *Ifnar1^fl/fl^*and *Ifnar1^fl/fl^ Lyz2-Cre* littermate controls. Age and sex-matched male and female mice between 6 and 10 weeks of age were enrolled in experiments. Mice were housed under specific pathogen-free conditions. Animal studies were approved by the animal care committee of the University of Toronto in accordance with the Canadian Council on Animal care.

### Cell culture

B16F10 cells obtained from the American Type Culture Collection and B16F10.OVA (B16.OVA) was obtained from Dr. Mikael Karlsson, Karolinska Institute. B16.OVA.tomato was generated using the Sleeping Beauty transposon system (Addgene #60513). The MC-38 cell line was obtained from Dr Jean Gariépy at the Sunnybrook Research Institute, Toronto. The cell lines were maintained at 37°C with 5% CO2 in complete Dulbecco’s modified Eagle’s medium [10% heat-inactivated fetal bovine serum (FBS), streptomycin (100 mg/ml), and penicillin (100 mg/ml); Gibco].

### In vivo tumor experiments

Mice were anesthetized with 2-3% isoflurane and 10^6^ B16.OVA, 2×10^6^ B16.OVA.tomato or 0.5 x 10^6^ MC-38 cells were inoculated intradermally or subcutaneously on the right flank in 50 µl D-PBS. Where relevant, mice were randomized into groups with similar average tumor sizes on the day of treatment. When tumors reached an average volume of 100 mm^3^, 25 µg of ADU-S100 (MedChemExpress) was administered intratumorally in 50 µl D-PBS. An equal volume of DMSO dissolved in D-PBS was used as vehicle control. Tumor volumes were determined with calipers and the formula (length x width^2^) x 0.5. Mice were euthanized when tumors reached >1500 mm^3^ or were severely ulcerated. For CCR5 blocking experiments, B16.OVA.tomato tumor-bearing C57BL/6 mice were given 10mg/kg maraviroc or PBS treatment 4 hours prior to ADU-S100 or vehicle treatment. LNs were collected for flow cytometry on day 1 post-treatment.

### Tissue processing

Tumors were resected and cut into small pieces and digested in 1 ml HBSS with 825 U/ml collagenase IV (Invitrogen) and 0.1 mg/ml DNAse I for 30 min at 37°C 250 rpm. Tumor cell suspensions were mashed through a 70 µM strainer, washed once in RPMI 10% FBS and resuspended at 100 mg of tumor/ml in D-PBS. Tumor-draining inguinal LNs on the same side as the tumor were mechanically disrupted through a 70 µM strainer to obtain single-cell suspensions.

### Flow cytometry

For flow cytometry, 2 x 10^6^ cells from the LN or 25 mg of tumor were used per stain. Cells were stained for 15 minutes in eF506 Viability Dye diluted in D-PBS, washed, and treated with Fc-receptor blocking antibodies anti-CD16/CD32 for 10 minutes (Invitrogen). Cells were then stained with cell surface antibodies for 30 minutes, washed and fixed for 10 minutes in Cytofix buffer (BD) at 4°C. For tetramer staining, cells were stained for 30 minutes with H-2K^b^/SIINFEKL-APC tetramers at room temperature before carrying out cell surface staining as described above. Anti-CD8a clone KT15 was used when staining with tetramers to prevent non-specific binding of H-2K^b^ tetramers that can occur with clone 53-6.7. For intracellular staining of nuclear proteins, cells were fixed and permeabilized with Foxp3 transcription factor staining buffer (Invitrogen) for 40 minutes prior to intracellular staining for 30 minutes at 4°C. For intracellular cytokine staining, cells were re-stimulated with 0.1 µM Trp-2 peptide for 6 hours prior to surface staining, then were fixed and permeabilized with Cytofix/Cytoperm buffer (BD) for 20 minutes prior to intracellular cytokine staining for 30 minutes at 4°C. Data were acquired on a LSR Fortessa X-20 or FACSymphony A3 using FACSDiva software (BD Biosciences) and analyzed with FlowJo (TreeStar). Absolute cell numbers in samples were determined using 123count eBeads (Invitrogen).

### Ifnar1 mixed bone marrow chimeras

CD45.1/.2 mice were irradiated with two doses of 550 rad with a 4h rest between doses, and the bone marrow was reconstituted by intravenous injection of a 1:1 mixture of *Ifnar1***^+^***^/^***^+^**CD45.1 and *Ifnar1^-/-^* CD45.2 bone marrow cells for a total of 5×10^6^ cells/mouse. Mice were fed neomycin sulfate water (2 mg/ml) for two weeks following irradiation. Chimerism was assessed in the blood >90 days later, prior to tumor cell inoculation.

### Sample preparation for CITE-sequencing

*Ifnar1* mixed bone marrow chimeras were inoculated with 10^6^ B16.OVA cells intradermally and a single dose of ADU-S100 was administered intratumorally on day 12. LNs and tumors were harvested at day 1 and day 2 post-treatment, respectively. LNs were processed as described above. Tumors were digested in HBSS 875 U/ml collagenase IV for 20 min at 37°C 250 rpm, filtered through 70 µM strainers, centrifugated and resuspended at 120 mg/ml in D-PBS. For both the tumor and LN, equal amounts of cell suspension from 3 vehicle or 3 ADU-S100-treated mice were pooled into one vehicle sample and one ADU-S100 sample. Cells were stained with eF506 fixable viability dye in PBS for 10 min on ice, then washed. Fc Receptors were blocked for 10 min before staining with fluor-conjugated antibodies (for FACS) and TotalSeq-A antibodies (Biolegend) in D-PBS 2% FBS for 25 min on ice. Here, vehicle and ADU-S100 treated samples were labeled with distinct hashtag antibodies, while anti-CD45.1 and anti-CD45 barcoded antibodies were used to label *Ifnar1***^+^***^/^***^+^**and *Ifnar1^-/-^* cells. Cells were washed three times with D-PBS and sorted on a BD FACS Aria IIu (tumor: live CD45**^+^** cells; LN: live CD3-CD19-CD11b**^+^**and/or MHCII**^+^** cells). Vehicle and ADU-S100-treated samples were pooled together at a 1:1 ratio immediately before partitioning on the 10X chromium controller. The Chromium Single Cell 3’ Reagent Kits v3.1 were used according to the manufacturer’s user guide in conjunction with CITE-seq & Cell Hashing Protocol from the New York Genome Center Technology Innovation Lab. Of note, anti-CD45 clone I3/2.3, anti-CD45 clone 30-F11 (in hashtag antibodies), anti-CD45.1 (clone A20) and anti-CD45.2 (clone 104) did not significantly interfere with each other under our experimental conditions.

### CITE-sequencing analysis

Sequencing reads from the mRNA library were processed through the Cell Ranger Single Cell Software Suite (10x Genomics) to obtain gene-cell expression matrices. Antibody-derived tags and hashtags were counted using CITE-seq count v1.4.2 and loaded into the Seurat object. RNA, ADT, and HTO expression matrices were imported into RStudio, and downstream analysis was performed using Seurat v3. Cells that expressed fewer than 500 or more than 7000 genes and had a percentage of mitochondrial genes over 10% were filtered out. RNA counts were log-normalized, while antibody-derived counts were normalized using centered log ratio transformation. The 2000 most variable genes were used for dimensionality reduction using principal component analysis. An elbow plot was used to identify the principal components that contribute to most of the variation. For the LN sample, we used the FindNeighbors function using 10 dimensions and ran the FindClusters function with a resolution of 0.5. For the tumor sample, we used the FindNeighbors function using 15 dimensions and ran the FindClusters function with a resolution of 0.5. Contaminating T cell, B cell, pDC, NK cell and neutrophil clusters were excluded from the samples and remaining cells were used for analysis. Differential gene expression analysis was performed in Seurat. Genes were considered significantly differentially expressed when they had a log_2_ fold-change of > 0.25 and an adjusted p. value < 0.05. To identify enriched gene sets, genes that were differentially expressed across monocyte/macrophage clusters were used as an input for overrepresentation analysis in g: Profiler using Gene Ontology: Biological processes gene sets. The SCENIC workflow was used to predict gene regulatory networks (27). NicheNet was used to predict receptor ligand interactions between cell types (28).

### In vivo IL-18 blockade

Mice were administered 200 µg of anti-IL-18 (clone YIGIF74-IG7, BioXCell) or isotype control (clone 2A3, BioXCell) in 100 µl D-PBS intraperitoneally one day before and one day after STING agonist administration.

### Statistical analysis

Data were analyzed in GraphPad Prism software (v9.3.1). Student’s t-test or one-way ANOVA were used to determine statistical significance as indicated in figure legends. Tumor growth curves were compared using two-way ANOVA with Sidak’s multiple comparisons test; the p-value at the last time point is reported. *P < 0.05, **P < 0.01, ***P < 0.001, and ****P < 0.0001 were applied.

### Data availability statement

Sequencing data will be made available on gene expression omnibus (GEO). Other data such as flow cytometry FCS files and tumor growth data are available from the authors upon request.

## Results

### STING agonist treatment induces distinct temporal peaks of MC accumulation in tumors and lymph nodes, respectively

Changes in myeloid cell populations in the tumors and tumor-draining LNs of mice were analyzed by flow cytometry following intra-tumoral injection of the STING agonist ADU-S100 into pre-established intradermal B16F10.OVA tumors (**Figure 1A**). One day post-treatment, the number and percentage of migratory cDC1 (CD26**^+^**MHCII**^hi^**CD11c**^+^**XCR1**^+^**), migratory cDC2 (CD26**^+^**MHCII**^hi^**CD11c**^+^**CD172**^+^**) and Ly6C^hi^ MCs (CD26^-^CD11b**^+^**Ly6G^-^Ly6C^hi^) was significantly increased in the LNs but decreased in the tumors (**Figure 1 B-D, F-G**). However, the number of Ly6C^hi^ MCs in the tumors was restored to initial levels by day 2 post-treatment (**Figure 1F**). Ly6C^hi^ MCs in the LN and tumor were CD26^-^CD88**^+^** whereas cDCs were CD26^hi^CD88^-^ (**Figure S1A, B**), supporting that these are monocytic lineage cells rather than inflammatory cDCs (29).

**Figure 1.**
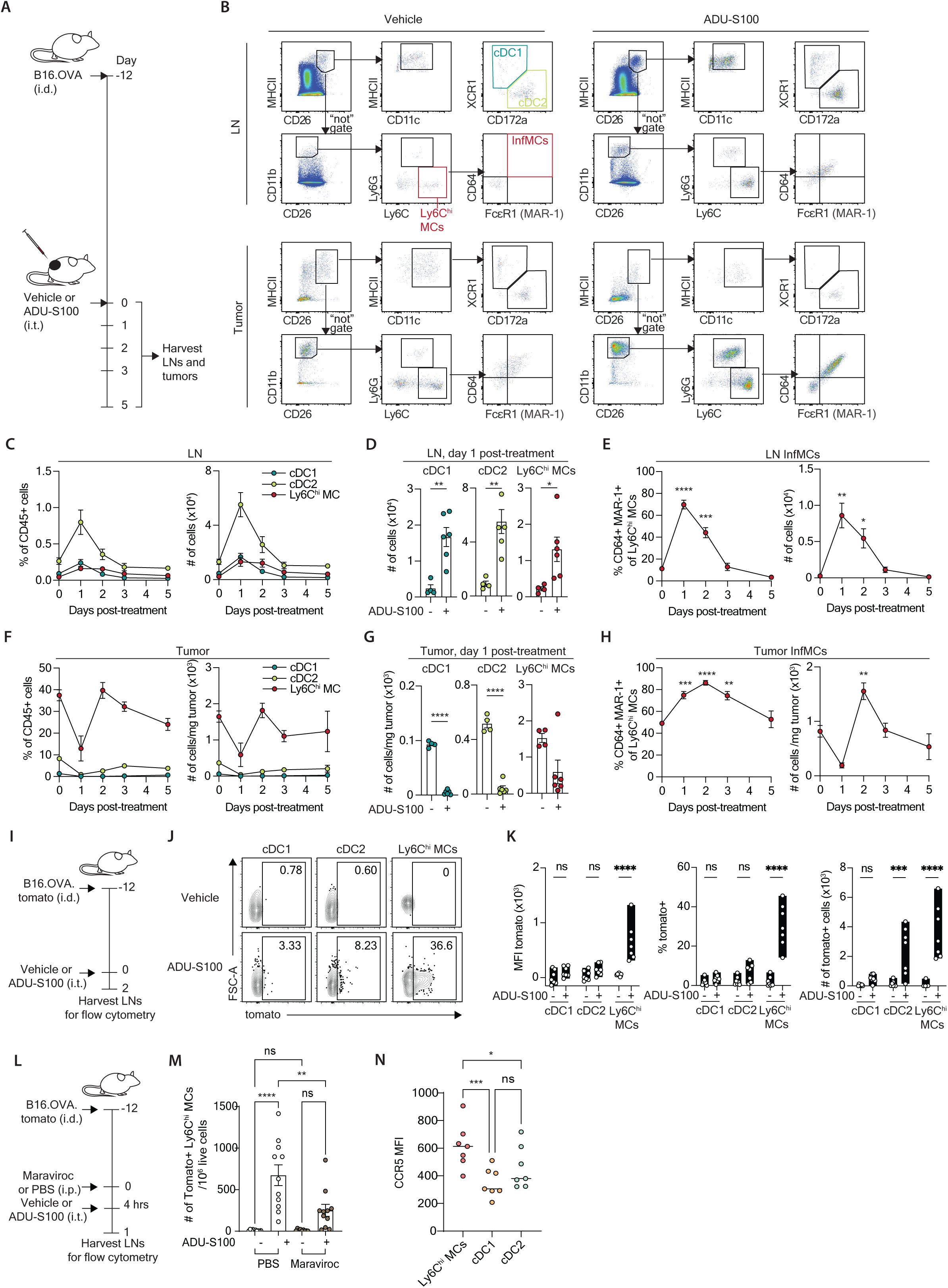
InfMCs accumulate in tumors and show CCR5-dependent trafficking to LN following STING agonist treatment. **(A-H)** STING agonists drive the accumulation of CD64^+^ MAR-1^+^ infMCs in the LN and tumor. **(A)** Experimental layout for panels (B-H). Mice were inoculated with 10^6^ B16.OVA cells intradermally and given an intratumoral injection of ADU-S100 or vehicle 12 days later. Tumors and draining LNs were harvested for flow cytometric analysis at 0-5 days post-treatment. (**B)** Gating strategy for identification of cDC1 (CD26**^+^**MHCII^hi^CD11c**^+^**XCR1**^+^**), cDC2 (CD26**^+^**MHCII^hi^CD11c**^+^**CD172**^+^**), Ly6C^hi^ MCs (CD26-CD11b**^+^**Ly6G-Ly6C**^+^**) and infMCs (CD26-CD11b**^+^**Ly6G-Ly6C**^+^**CD64**^+^**MAR-1**^+^**) in the LN and tumor, starting from single, live CD45**^+^** cells. **(C-H)** Number and percentage of cells in the LN (C-E) and tumor (F-H) at indicated time points after treatment with ADU-S100 or vehicle. **(I-K)** InfMCs carry tumor antigens to the LN. **(I)** Experimental layout for panels (J,K). C57BL/6 mice were inoculated with 10^6^ B16.OVA.tomato cells intradermally and given ADU-S100 or vehicle 12 days later. LNs were collected for flow cytometry on day 2 post-treatment. **(J)** Representative flow plots. **(D)** Tomato signal in LN APC populations depicted as the tomato MFI (left panel), % tomato**^+^** (middle panel) and absolute number of tomato**^+^** cells (right panel). (L) Experimental design for panels (M-N). B16.OVA.tomato tumor-bearing C57BL/6 mice were given 10mg/kg maraviroc or PBS treatment 4 hours prior to ADU-S100 or vehicle treatment. TDLNs were collected for flow cytometry on day 1 post-treatment. (M) Tomato signal in LN Ly6C^hi^ MC population depicted as absolute number of tomato^+^ cells. (N) CCR5 expression on Ly6C^hi^ MCs, cDC1 and cDC2 in TDLN one day after ADU-S100 treatment. Data are pooled from two or three independent experiments, each with 3-6 mice per group. Unpaired t-test (D,E,G,H), two-way ANOVA (K,M), or one-way ANOVA with Sidak’s multiple comparisons test (N) was performed.

CD64 and MAR-1 staining can be used to identify infMCs in mice, as these antibodies label cells that have responded to inflammatory cytokines and PAMPS (29). While CD64 is an Fc receptor upregulated by monocytes in response to IFN-Is and other pro-inflammatory stimuli, MAR-1 is an antibody clone raised against FcεR1 but that was later found to bind to other Fc receptors including CD64 (29, 30). Here, we used these markers to identify cells that had potentially responded to IFN-Is downstream of intra-tumoral STING agonist administration. We also noted that both the cDC and the Ly6C^hi^ MCs had higher levels of MHC I after ADU-S100 compared to vehicle control, consistent with an IFN-I dependent response (**Figure S1C**). The number and percentage of Ly6C^hi^ MCs positive for CD64 and MAR-1 were significantly increased in the LNs and tumors following STING agonist therapy and peaked at 1- and 2-days post-treatment, respectively (**Figure 1E, H**). Thus, despite an initial decrease in the number of Ly6C^hi^ MCs in the tumor at day 1 post-treatment, the absolute number of infMCs in tumors was increased by day 2 post-treatment compared to baseline. Taken together, these data show that intra-tumoral administration of ADU-S100 drives the accumulation of CD64**^+^** MAR-1**^+^** infMCs in the LNs and tumors.

The ability of Ly6C^hi^ MCs to migrate from tissues to the LN and cross-present antigens during viral infection has been questioned (29, 31). To test if the significantly increased number of Ly6C^hi^ MCs in the LN on day 1 following STING agonist therapy was due to migration from the tumors, wild-type mice were inoculated with B16F10.OVA (B16.OVA) cells that express the red fluorescent protein TdTomato (B16.OVA.tomato) and treated with ADU-S100 or vehicle (**Figure 1I**). In these mice, intra-tumoral STING agonist injection led to a substantial increase in the number and percentage of tomato**^+^** CD26^-^CD11b**^+^**Ly6G-Ly6C^hi^ MCs in the LN (**Figure 1J-K**). We also observed a smaller increase in the number and percentage of tomato**^+^** cDC2 (**Figure 1J-K**).

Maraviroc is a CCR5 antagonist that has been shown to block CCR5-dependent monocyte trafficking to the LNs (32). Therefore, to determine if CCR5 plays a role in trafficking of the tomato**^+^**CD26^-^CD11b**^+^**Ly6G-Ly6C^hi^ MCs from the tumor to the LN, we treated wild-type mice bearing B16.OVA.tomato tumor with maraviroc or vehicle control, then 4 hours later with ADU-S100 or vehicle and monitored tomato^+^ in the tumor dLN at 24 hours (**Figure 1L**). Consistent with a role for CCR5 in migration of monocytes to LNs, maraviroc treatment substantially reduced accumulation of the tumor antigen bearing monocytes to the dLN (**Figure 1M**). We also noted that LN monocytes from STING agonist treated mice express higher CCR5 than cDC1 and cDC2 at 1 day after STING agonist treatment (**Figure 1N**). Taken together, these data suggest that STING agonist treatment induced accumulation of monocytic lineage cells in LN and tumor at different time points, and monocytic lineage cells carry tumor protein to the tumor-draining LN in a CCR5-dependent manner.

### infMCs are required for optimal tumor control following intra-tumoral STING agonist injection

CCR2 signaling is required for exit of monocytes from the bone marrow (33, 34). To test if CCR2-dependent infMCs play a role in the tumor control induced by STING agonists, we tracked the growth of B16.OVA and MC-38 tumors in *Ccr2***^+^***^/^***^+^** and *Ccr2^-/-^* littermate mice treated with ADU-S100 or vehicle control. In contrast to *Ccr2***^+^***^/^***^+^**mice, infMCs do not accumulate in the tumors of *Ccr2^-/-^* mice after STING agonist therapy (**Figure 2A**), whereas tumor accumulation of other immune cell types was unaffected (**Figure S2A, B**). While *Ccr2***^+^***^/^***^+^**and *Ccr2^-/-^* mice treated with the vehicle control showed no differences in tumor growth, *Ccr2^-/-^*mice showed impaired tumor control compared to *Ccr2***^+^***^/^***^+^**mice following STING agonist therapy (**Figure 2B, C**).

**Figure 2.**
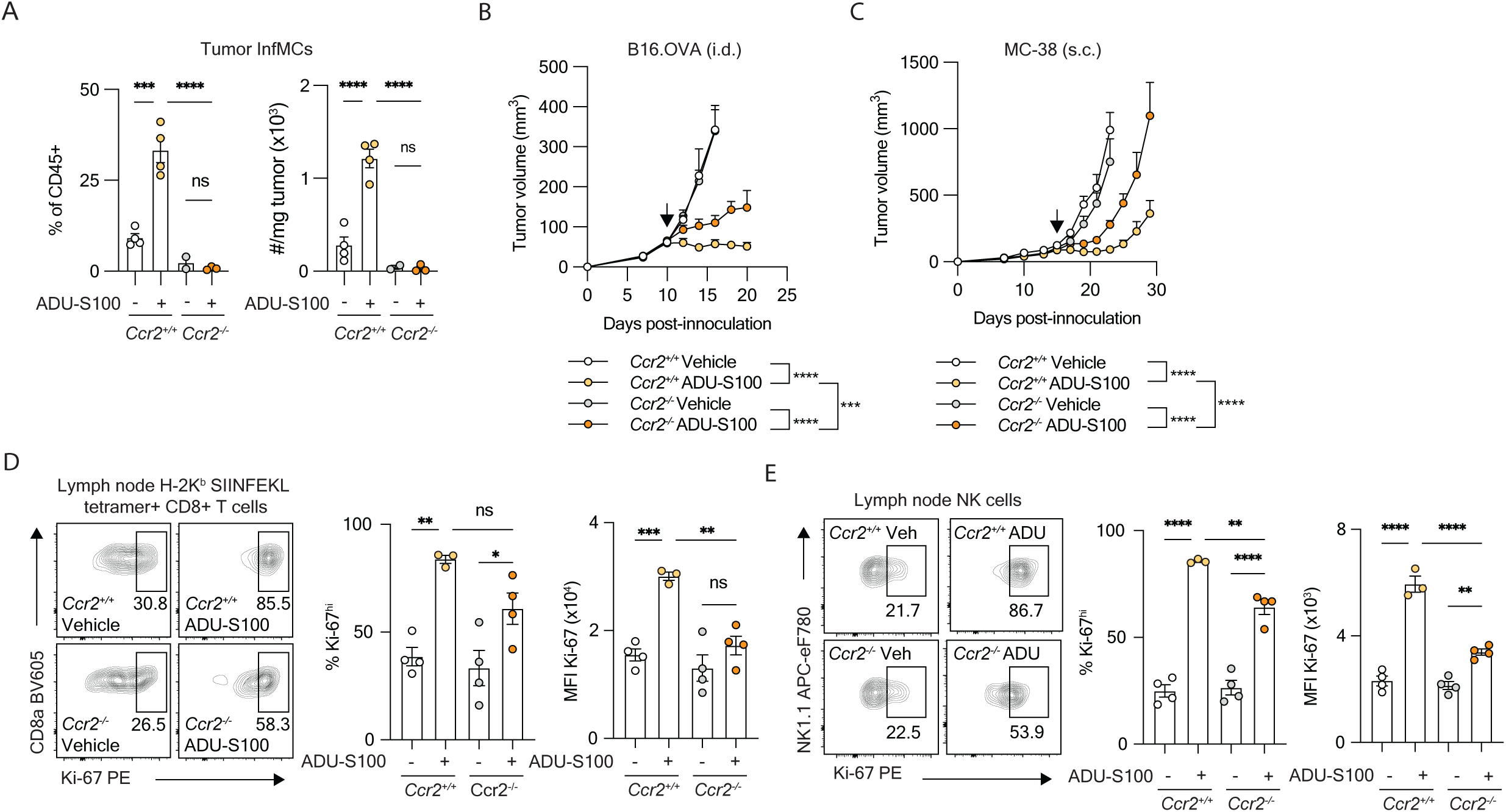
InfMCs are required for optimal responses to STING agonist therapy. **(A)** Number and percentage of infMCs in B16.OVA tumors of *Ccr2***^+^***^/^***^+^** and *Ccr2^-/-^* mice two days post-treatment with ADU-S100 or vehicle. **(B-C),** Volume of B16.OVA intradermal tumors (B) or MC-38 subcutaneous tumors (C) in *Ccr2***^+^***^/^***^+^** and *Ccr2^-/-^* mice treated with ADU-S100 or vehicle on days indicated by an arrow. **(D-E),** Ki-67 staining on LN H-2K^b^ SIINFEKL tetramer**^+^** CD8**^+^** T cells (D) or NK cells (E) three days post-treatment with ADU-S100 or vehicle. Data are representative of two independent experiments. One-way ANOVA (A, D-E), or two-way ANOVA with Sidak’s multiple comparisons test (B,C) was performed.

STING agonist therapy induces the expansion of tumor specific CD8**^+^**T cell and NK cell populations which are measurable as early as 3 days post-treatment (18, 21). To test if monocytes contribute to the proliferation of tumor-specific T cells and NK cells, *Ccr2***^+^***^/^***^+^** and *Ccr2^-/-^* mice with B16.OVA tumors were sacrificed 3 days post-treatment with STING agonists or vehicle control for flow cytometry analysis. STING agonists significantly increased the proportion of H-2K^b^ SIINFEKL**^+^** CD8**^+^** T cells and NK cells proliferating in the LN based on Ki-67 staining (**Figure 2D, E and Figure S2C**). *Ccr2^-/-^* mice showed reduced Ki67^+^ NK cells (% and mean fluorescence intensity, MFI), but there was no impact on the absolute number of Ki67^+^ NK cells in LN and tumor (**Figure 2E and S2C**). For T cells, although the MFI for Ki67 was decreased in CCR2^-/-^ mice, neither the frequency or absolute number of Ki67^+^ cells was significantly impacted by CCR2 (**Figure 2D and S2C**). These results may be confounded by the significant apoptosis of lymphocytes in tumors early after ADU-S100 treatment (35). At later time points rapid tumor shrinkage in the *Ccr2^+/+^* mice precluded accurate T cell measurements. Therefore, to further investigate whether *Ccr2*-deficiency affects early T cell proliferation, we adoptively transferred OT-I T cells into *Ccr2^+/+^* or *Ccr2^-/-^* B16.OVA tumor bearing mice 1 day prior to ADU-S100 treatment. However, there was no significant impact of *Ccr2* deficiency on LN OT-I T cell proliferation (**Figure S2D**). These data suggest that CCR2-dependent MCs contribute to tumor control but have little or no impact on the initial proliferation of the tumor-specific T cells, suggesting that MCs do not contribute to cross-priming in this model.

### InfMCs are driven by intrinsic IFN-I signalling during STING agonist therapy

Mice deficient for the interferon alpha/beta receptor 1 (IFNAR1) have sub-optimal responses to STING agonists (16, 18), but whether IFNAR1 signaling within the monocytes promotes anti-tumor immunity in this context is unknown. To address this question, we tracked the growth of B16.OVA tumors in *Ifnar1^fl/fl^ Lyz2-Cre* mice, which have lower expression of IFNAR1 on myeloid cells, including monocytes (**Figure 3A, B**). *Ifnar1^fl/fl^ Lyz2-Cre* mice showed increased tumor growth following STING agonist treatment compared to ADU-S100 *Ifnar1^fl/fl^* littermate controls (**Figure 3C**). These data suggest that IFN-I responses in infMCs are required for optimal tumor control in the presence of STING agonists.

**Figure 3.**
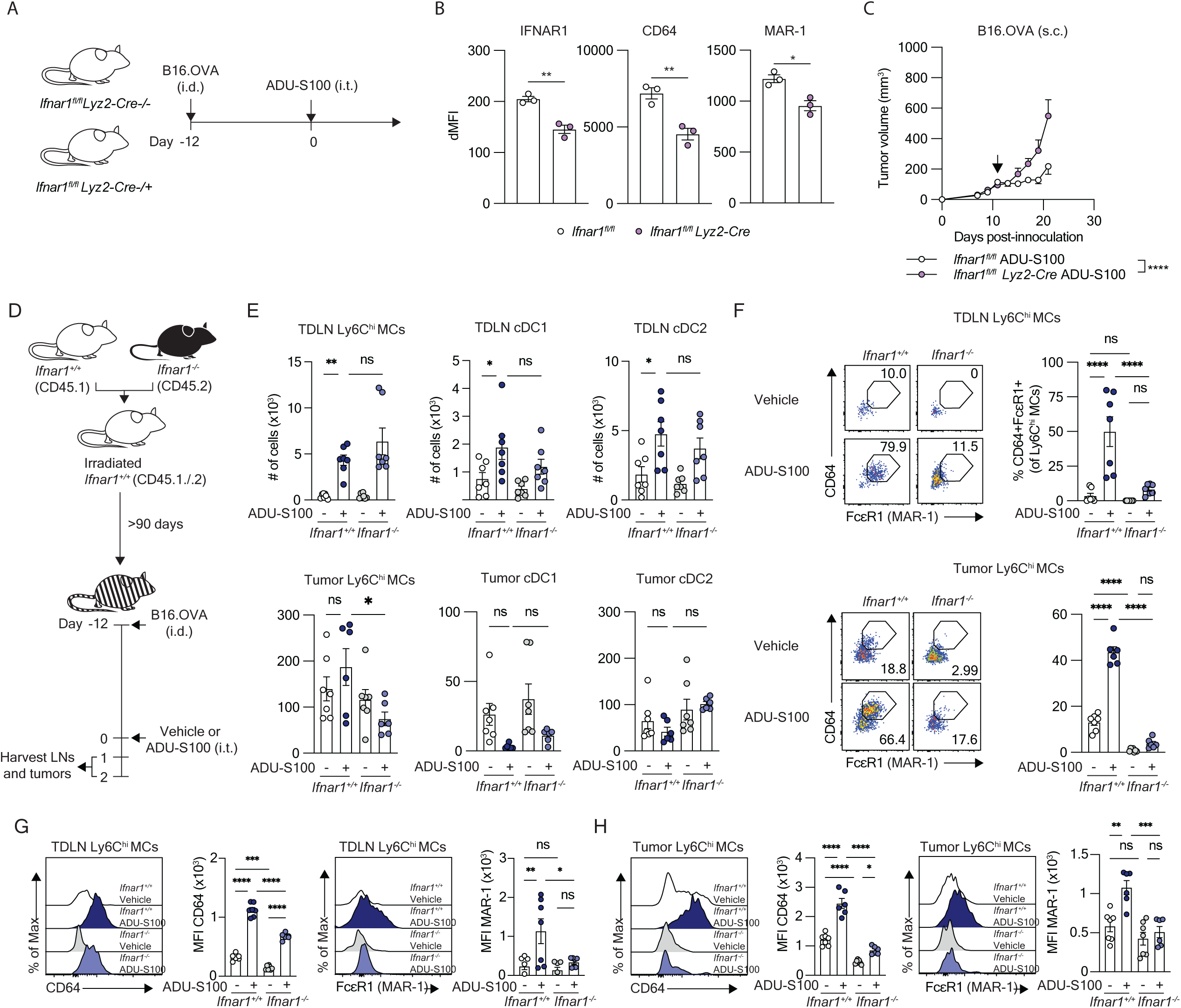
Intrinsic IFN-I signaling drives an inflammatory phenotype in Ly6C^hi^ MCs but is dispensable for their migration to the LN. **(A)** Experimental layout for panels (B,C). *Ifnar1^fl/fl^* and *Ifnar1^fl/fl^ Lyz2-Cre* mice were inoculated with B16.OVA tumors and treated with ADU-S100 or vehicle 12 days later. **(B)** IFNAR1, CD64 and MAR-1 staining on Ly6C^hi^ MCs from the B16F10.OVA tumors of *Ifnar1^fl/fl^*and *Ifnar1^fl/fl^ Lyz2-Cre* mice on day 5 post-treatment. **(C)** Volume of B16F10.OVA tumors in *Ifnar1^fl/fl^* and *Ifnar1^fl/fl^ Lyz2-Cre* mice treated with ADU-S100 on day 11. **(D)** Experimental layout for panels (E-G). Irradiated CD45.1/.2 mice were reconstituted with a 1:1 mixture of CD45.1 *Ifnar1*^+*/*+^ and CD45.2 *Ifnar1^-/-^* bone marrow cells. After >90 days reconstitution, mice were inoculated with 10^6^ B16F10.OVA cells intradermally and given ADU-S100 or vehicle intratumorally 12 days later. TDLNs and tumors were collected 1- and 2-days post-treatment, respectively for flow cytometric analysis. **(E)** Absolute numbers of *Ifnar1*^+*/*+^ and *Ifnar1^-/-^* cDC1, cDC2 and Ly6C^hi^ MCs in the TDLNs (top) and tumors (bottom) of the mixed bone marrow chimeric mice. **(F)** Percentage of Ly6C^hi^ MCs that express the infMC markers CD64 and MAR-1 in the TDLNs (top) and tumors (bottom) of the mixed bone marrow chimeric mice. **(G)** Mean fluorescent intensity of CD64 and MAR-1 on Ly6C^hi^ MCs in the TDLNs (top) and tumors (bottom) of the mixed bone marrow chimeric mice. Data are pooled from two independent experiments with 3-4 mice each.Unpaired t-test (B), two-way ANOVA with Sidak’s multiple comparisons test (C), or One-way ANOVA with Tukey’s multiple comparisons test (E-G) was performed.

To more directly test if infMCs were driven by IFN-I, we analysed responses to STING agonist therapy in mixed bone marrow chimeras in which the hematopoietic cells were derived from a 1:1 mix of *Ifnar1***^+^***^/^***^+^**or *Ifnar1^-/-^* bone marrow cells (**Figure 3D**). The increase in numbers of cDC1, cDC2 and Ly6C^hi^ MCs in the LN at day 1 following STING agonist therapy was largely independent of intrinsic IFN-I signaling (**Figure 3E**). However, upregulation of inflammatory markers CD64 and MAR-1 by Ly6C^hi^ MCs in the LN and tumor was mainly restricted to the wild-type compartment of the chimeric mice (**Figure 3F-H**). These data indicate that intrinsic IFN-I signaling drives an inflammatory phenotype in Ly6C^hi^ MCs but is not required for MC migration to the LN during STING agonist therapy.

To define how intrinsic IFN-I signalling downstream of intratumoral STING agonist treatment alters the transcriptional state of myeloid cells, CITE-sequencing was performed on immune cells derived from the LNs and tumors of *Ifnar1* mixed bone marrow chimeric mice treated with ADU-S100 or vehicle control (**Figure 4A**). Mice were reconstituted with a 1:1 mix of *Ifnar1***^+^***^/^***^+^**and *Ifnar1^-/-^* bone marrow cells, and chimerism in the blood was confirmed after 90 days later (**Figure S3A, B**). At 12 days following tumor inoculation, mice were treated with ADU-S100 or vehicle and LNs and tumors were harvested 1 day and 2 days post-treatment, respectively. To enrich for myeloid cells, CD3-CD19-CD11b**^+^** and/or MHCII**^+^** cells were sorted from the LN. Because the tumors are already enriched for myeloid cells, total CD45**^+^** cells were sorted from tumors (**Figure S3C).** Here, DNA-barcoded antibodies were used to trace the treatment group (vehicle or ADU-S100) and the genotype (*Ifnar1***^+^***^/^***^+^**or *Ifnar1^-/-^*) of each single cell based on the sequencing data (**Figure S3D, E**). After quality filtering, 12, 078 and 3001 cells from the LN and tumor, respectively, remained for downstream analysis (**Figure S3F**). As expected, cells assigned as *Ifnar1^-/-^* cells expressed very low levels of ISGs compared to *Ifnar1***^+^***^/^***^+^**, and ISG expression levels were higher in cells derived from the ADU-S100-treated group compared to vehicle, validating the demultiplexing strategy (**Figure S3G**).

**Figure 4.**
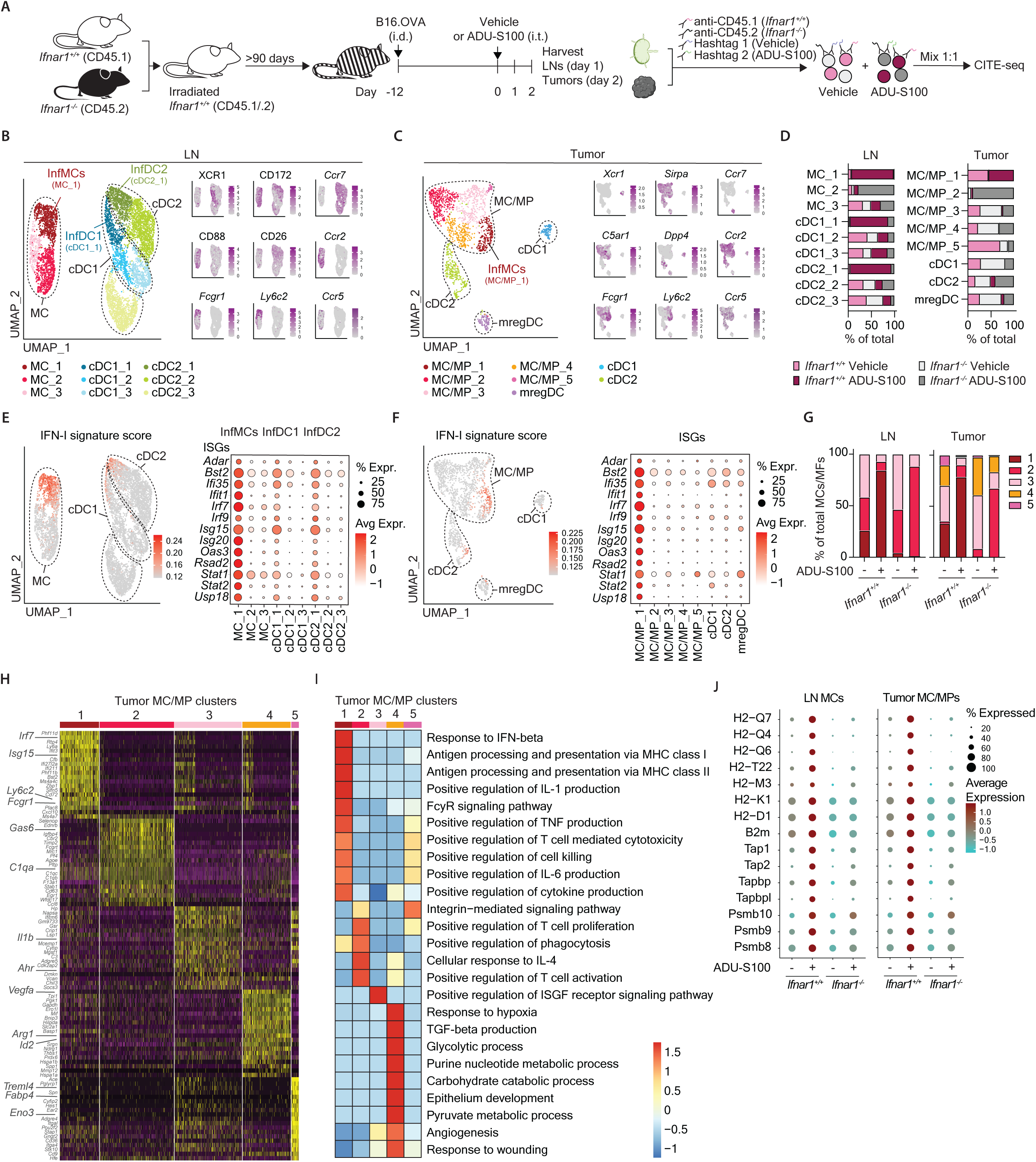
Intrinsic IFN-I signaling drives an inflammatory transcriptional state in LN and tumor monocytes. CITE-sequencing analysis of myeloid cells from the LNs and tumors of *Ifnar1* mixed bone marrow chimeric mice on days 1 and 2 post-treatment with ADU-S100 or vehicle. **(A)** Experimental design. **(B-C)** UMAP of cDC, monocyte and macrophage populations in the LN. (B) and tumor (C). **(D)** Proportion of cells in each cluster derived from each genotype and treatment condition. **(E-F)** UMAP depicting the IFN-I signature score for each cell in the LN (E) and tumor (F). On the right, dotplots depicting the expression of common ISGs across clusters. **(G)** Percentage of total monocytes/macrophages in each group derived from each cluster. **(H)** Heatmaps depicting top DEGs across or tumor monocyte/macrophage clusters. **(I)** Pathway enrichment analysis (g:Profiler, GO biological processes) using DEGs for each tumor monocyte/macrophage cluster (Mono/Mac 1-5). **(J)** Expression of genes involved in antigen processing and presentation via MHC class I in monocyte/macrophages from the LN and tumor of *Ifnar1* mixed bone marrow chimeric mice treated with ADU-S100 or vehicle.

Clusters of monocytes/macrophages (MC/MP) (CD88/*C5ar1*, CD11b/*Itgam*), cDC1 (XCR1/*Xcr1*, CD26/*Dpp4*) and cDC2 (CD172/*Sirpa*, CD26/*Dpp4*) were identified in the LN and tumor (**Figure 4B-C**). LN cDCs also expressed high levels of *Ccr7*, as opposed to monocytes, which expressed *Ccr2* and *Ccr5* (**Figure 4B-C**). Each cell type could be further subdivided into subclusters that represent different transcriptional states. Clusters MC_1, cDC1_1 and cDC2_1 in the LN and cluster MC/MF_1 in the tumor were almost exclusively derived from the *Ifnar1***^+^***^/^***^+^**compartment of the STING agonist treated chimeric mice (**Figure 4D**). These clusters shared a common IFN-I gene signature with high expression of interferon signaling genes (ISGs) including *Isg15*, *Isg20*, *Irf7*, *Ifit1*, *Ifi35* and *Bst2* (**Figure 4E-F**). The proportion of MC/MPs with this inflammatory signature was substantially higher in the STING agonist-treated mice compared to controls (**Figure 4G**). Moreover, this state was entirely dependent on intrinsic IFN-I signalling as no *Ifnar1^-/-^* cells were found in these clusters (**Figure 4G**). Because clusters MC_1, cDC1_1 and cDC2_1 in the LN and MC/MP_1 in the tumor are driven by IFN-Is, we will refer to them as infMCs, infDC1 and infDC2.

In the LN and tumor, cells in cluster MC_2 were almost uniquely derived from the *Ifnar1*^-/-^ compartment of STING agonist-treated mice, suggesting this state is induced by STING agonist therapy in the absence of intrinsic IFN-I signalling (**Figure 4D**). Cells from clusters MC/MP_3 and MC/MP_4 in the tumor were derived from both treatment groups and genotypes, indicating these states are not driven by STING agonist therapy or IFN-I signaling (**Figure 4D**). Cluster MC/MP_4 expressed genes associated with pro-tumorigenic TAMs such as *Vegfa*, *Arg1* and *Cd274* (**Figure 4H**). Pathways associated with cells in tumor cluster MC/MP_4 include “Angiogenesis” and “TGF-β production”, suggesting they play a pro-tumorigenic role (**Figure 4I**). The proportion of MC/MP4s was reduced upon STING agonist therapy in an IFN-I-dependent manner (**Figure 4G**). LN and tumor monocyte derived cells also showed IFNAR-1 dependent increases in MHC I genes, along with TAP and proteasome genes (**Figure 4J**). Taken together, these data show that intrinsic IFN-I signaling reduces pro-tumorigenic gene signatures while enhancing genes associated with antigen presentation and inflammatory cytokine production in monocyte-derived cells.

### InfMCs have a unique inflammatory gene signature compared to infDCs

Inflammatory gene signatures of infMCs, infDC1 and infDC2 were determined by differential gene expression analysis between inflammatory and non-inflammatory subclusters. A core set of 214 inflammatory genes were shared between infMCs, infDC1 and infDC2 (**Figure 5A**). These included common ISGs such as *Isg15*, *Isg20* and *Ifit1*. Inflammatory genes that were more highly expressed by infMCs included the cytokine *Il18*, the chemokines *Ccl2*, *Cxcl9* and *Cxcl10*, cell surface molecules *Fcgr1* and *Ly6c2*. In contrast, infDCs expressed higher levels of *Il15*, *Cd70*, *CD86* and *Cd274* (**Figure 5B**). Regulatory network inference predicted that the inflammatory programs in infMCs, infDC1 and infDC2 are driven by different transcriptional regulators downstream of IFN-I signaling. While infMCs were driven by ISGF3 (*Stat1*, *Stat2* and *Irf9*), *Irf5* and *Irf7*, infDCs were predominantly driven by NF-κB family members (*Nfkb2*, *Rel*, *Relb*), *Stat3*, *Stat4* and *Klf6* (**Figure 5C**).

**Figure 5.**
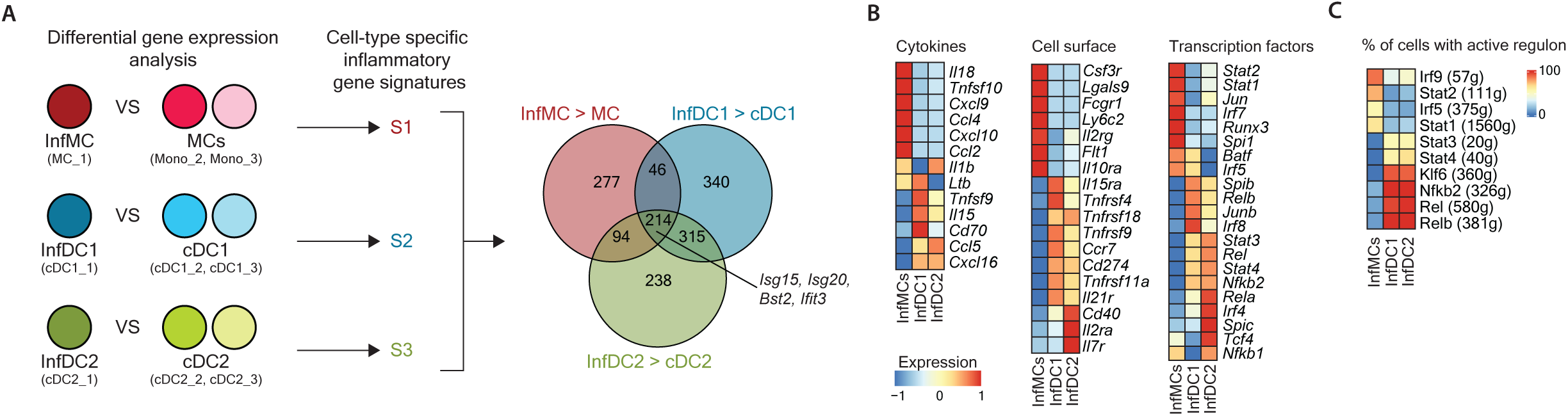
InfMCs have a unique inflammatory program compared to infDCs. **(A)** Inflammatory signatures for infMCs, infDC1 and infDC2 were calculated and compared in a Venn diagram. **(B)** Heatmaps depicting differential expression of selected inflammatory genes across infAPC populations. **(C)** Heatmap depicting the percentage of cells in each infAPC population in which the indicated regulons are active, as determined by single-cell regulatory network inference analysis.

By comparing the changes in gene expression between monocytes and macrophages from vehicle and STING-agonist treated mice within the *Ifnar1*^+/+^ and *Ifnar1*^-/-^ compartments, we identified a set of genes that were upregulated upon STING agonist therapy in both the *Ifnar1^+/+^* and *Ifnar1^-/-^* compartments (**Figure S4**). These included genes involved in the complement system (*C1qa, C1qb, C1qc*, *C3ar1*), genes associated with lysosomes including cathepsins (*Ctsb, Ctsc, Ctsd, Ctsl*) and *Lamp1*, as well as genes encoding for the transcription factors *Fosb* and *Maf* (**Figure S4**).

### Il18 is induced by IFN-I signalling in monocytic lineage cells and is important for the anti-tumor response to STING agonists

Receptor-ligand interactions between infMCs and CD8**^+^**T cells or NK cells were predicted using NicheNet (28). IL-18 – IL-18R1 interaction was the top predicted interaction between infMCs and CD8**^+^** T cells or NK cells (**Figure 6A**). *Il18* expression was specifically upregulated by infMCs and not by infDC1 or infDC2 (**Figure 6A**). IL-18R1 was highly expressed on both tumor-specific CD8**^+^** T cells and NK cells in the lymph node and tumor and was upregulated on tumor-specific CD8**^+^** T cells upon STING agonist therapy (**Figure 6B-D**). Accordingly, blockade of IL-18 using a neutralizing antibody led to impaired tumor control in mice treated with ADU-S100 (**Figure 6E-F**), in line with previous studies that show that activation of the IL-18-IL-18R pathway promotes CD8**^+^** T cell and NK cell responses against tumors (36, 37). To determine the contribution of monocyte derived IL-18 to tumor control, we compared the effect of blocking IL-18 with anti-IL-18 antibody in *Ccr2^+/+^*or *Ccr2^-/-^* mice (**Figure 6G**). Whereas anti-IL-18 increased tumor growth in WT mice, tumor growth was not impacted by anti-IL-18 in CCR2^-/-^ mice consistent with a role for infMC in provision of IL-18. In addition, we found that blockade of IL-18 in STING agonist treated mice impaired IFN-γ production by tumor Ag-specific CD8 T cells in the LN (Figure 6H-J). These data support a role for monocyte-derived IL-18 in increasing CD8 T cell production of IFN-γ in the dLN, resulting in increased tumor control during STING agonist therapy.

**Figure 6.**
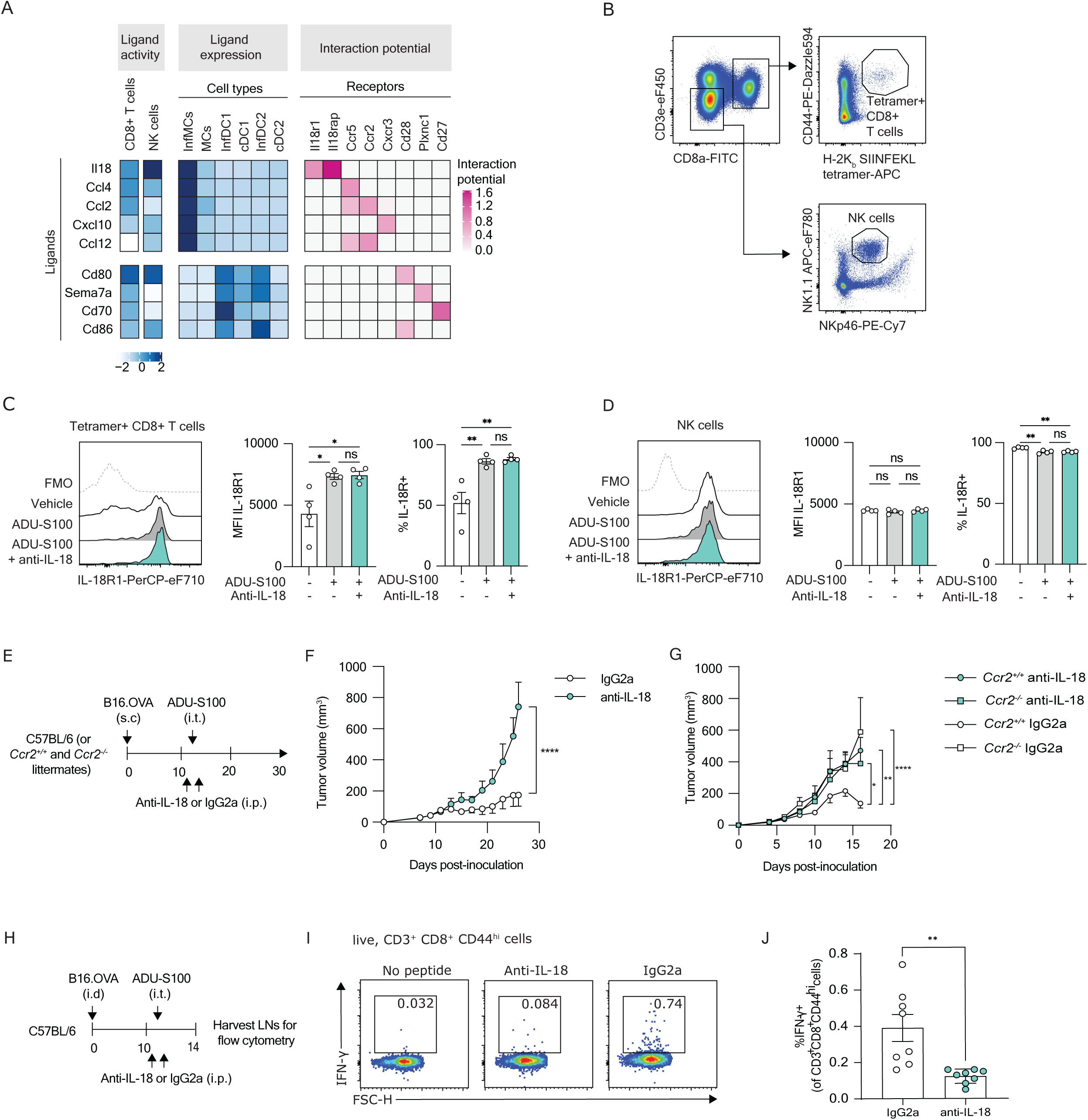
IL-18 is required for optimal tumor control in response to STING agonists. **(A)** NicheNet analysis of receptor ligand interactions between infAPC populations and CD8**^+^**T cells or NK cells. **(B)** Gating strategy to identify H-2K^b^ SIINFEKL tetramer**^+^** CD8**^+^** T cells and NK cells in the LNs starting from live CD45**^+^**cells. **(C-E)** Mice were administered 200 µg anti-IL-18 or control IgG2a intraperitoneally twice: one day before and one day after intra-tumoral administration of ADU-S100 or vehicle control. **(C-D)** Expression of IL-18R1 on LN H-2K^b^ SIINFEKL tetramer**^+^** CD8**^+^** T cells (C) and NK cells (D) 3 days post-treatment with ADU-S100. **(E)** Experimental design for panels (F-G)**. (F)** Volume of B16.OVA tumors in mice treated with anti-IL-18 or IgG2a and ADU-S100. (**G**) Volume of B16.OVA tumors in *Ccr2***^+^***^/^***^+^**and *Ccr2^-/-^* littermate mice treated with anti-IL-18 or IgG2a and ADU-S100. **(H)** Experimental design for panels (I-J) **(I)**. Representative flow cytometry data for IFN-γ production by live CD8 CD44^hi^ T cells in the dLN, day 3 after STING agonist treatment, showing no peptide control, as well as restimulation with Trp-2 peptide, for the anti-IL-18 or isotype control treated mice. **(J)**. Summary of two experiments with 3-4 mice per group per experiment. Data are representative of two independent experiments, each with 4-8 mice per group. One-way ANOVA (C-D), two-way ANOVA with Sidak’s multiple comparisons test (F-G), or unpaired t-test (J) was performed.

## DISCUSSION

MCs often acquire pro-tumorigenic features in the TME in response to factors such as hypoxia and fibrosis. Therefore, therapeutic strategies have aimed at either depleting these populations or reprogramming them towards more pro-immunogenic states (38). The latter approach may be preferable, as it takes advantage of the anti-tumorigenic features of MCs that are non-redundant with that of other immune cells. STING agonist treatment promotes recruitment of MCs, and the IFN-I production induced by STING agonists drives intrinsic phenotypical, transcriptional, and functional changes in the MC, associated with tumor control.

We show here that *Ccr2*^-/-^ mice have impaired responses to STING agonist therapy compared to *Ccr2***^+^***^/^***^+^** littermates. CCR2 is required for monocyte egress from the bone marrow, and *Ccr2*^-/-^ mice lack circulating monocytes (33, 34). Although we cannot exclude that *Ccr2*^-/-^ mice might also have defects in the CCR2-dependent migration of other cell populations, the finding that CCR2-deficiency did not impact overall numbers of cDC1 or cDC2, is consistent with monocytes playing a role in STING agonist therapy. Previous studies have reported reduced response to STING agonists in cancer models when mice are treated with chlodronate liposomes (23, 25), which deplete phagocytic cells including macrophages and MCs. Our results complement these studies by suggesting a more specific role of CCR2-dependent MCs, as opposed to tissue macrophages, during STING agonist therapy of cancer and by investigating the intrinsic role of IFN-I signaling in these cells.

Our experiments using *Ifnar1^fl/fl^ Lyz2-Cre* mice indicate that intrinsic IFN-I signaling in *Lyz2*^+^ cells contribute to tumor control in the presence of STING agonists. *Lyz2*-cre is expected to delete floxed genes in approximately 40-50% of blood monocytes with less than 10% deletion in cDCs (39). Accordingly, we observed a significant loss of cell surface IFNAR1 expression on tumor-infiltrating Ly6C^hi^ MCs in *Ifnar1^fl/fl^ Lyz2-Cre* compared to *Ifnar1^fl/fl^* controls. This loss of IFNAR expression was sufficient to significantly reduce cell surface expression of the IFN-I regulated protein CD64 on these cells, validating that this mouse model is relevant to study the role of IFN-I responses in Ly6C^hi^ MCs. While we cannot exclude that IFN-Is in other *Lyz2*^+^ cells such as neutrophils might also contribute to the response to STING agonist therapy, these data combined with the experiments using *Ccr2^-/-^* mice are compatible with a role for IFN-I signaling in monocytes during STING agonist therapy of cancer.

Some studies have shown a role for monocytes as a source of STING induced IFN-I. For example, Lam et al showed that microbiota-derived STING agonists induced by a high fat diet induced IFN-I production by tumor monocytes to promote DC-NK cell crosstalk (40). Monocytes were also recently found to be the main source of IFN-I after radiation therapy of mice (26), whereas another recent study showed that tumors are the major source of STING-dependent IFN-I in mouse and human cancers (41). We did not observe *Ifn* production by monocytes in our RNA-sequencing data at day 2 post-treatment, albeit we did not further pursue the source of IFN-I in our model.

A recent study by Tadepalli et al. used depletion of CCR2 positive cells during radiation therapy of mouse tumors to implicate monocytes in CD8 T cell responses and tumor control in a model in which STING is important for the effects of radiation induced tumor control (26). Conversely, Nicolai et al. used IFNAR1 CD11c-cre mice to show that IFN-I responses in cDCs are required for NK-cell mediated tumor control during STING agonist therapy (21). Our findings are complementary to these previous studies on the role of STING during tumor therapy because they highlight the role of CCR2-dependent cells and intrinsic IFN-I signaling in MCs following intratumoral delivery of STING agonists (18, 21, 26, 40, 41).

Using mixed bone marrow chimeric mice, in which half the hematopoietic cells lack IFNAR1, combined with flow cytometry and CITE-sequencing, we showed that intrinsic IFN-I signaling in monocytic lineage cells drives an inflammatory phenotype characterized by positive staining for CD64 and FcεR1 (MAR-1), akin to infMCs observed in models of acute viral infection (4, 5). Transcriptionally, infMCs express high levels of common ISGs, as well as inflammatory cytokines (*Il15*, *Il18*) and chemokines (*Ccl2, Cxcl9, Cxcl10*). This state is also entirely dependent on intrinsic IFN-I signaling, as *Ifnar1^-/-^*cells within the same mice do not adopt this transcriptional state, despite exposure to the same environmental cues. Although STING agonists can also induce other cytokines such as TNF and IL-6 (42), the monocyte response is heavily weighted toward IFNAR signaling with only a handful of IFNAR-independent STING induced genes observed (Fig S4). In contrast, the IFN gene signature in DC1 and 2, while present, is weaker than that of monocytes (Figure 4E). Inflammatory DC1, DC2 and infMC shared a core of 214 inflammatory genes. However, analysis of the single cell data by single-cell regulatory network inference analysis suggested while infMCs were driven by ISGF3 (*Stat1*, *Stat2* and *Irf9*), *Irf5* and *Irf7*, infDCs were predominantly driven by NF-κB family members (*Nfkb2*, *Rel*, *Relb*), *Stat3*, *Stat4* and *Klf6,* which is consistent with other cytokines such as TNF and IL-6 also playing a role. Although not further investigated here, this likely reflects the distinct epigenetic and transcriptional state of the different cell types, leading to distinct outcomes.

Skewing of monocyte and macrophage populations towards a more pro-inflammatory phenotype during STING agonist therapy was previously reported (23,25), but whether this was driven by intrinsic IFN-I responses had not been tested until now. In line with the study by Xu et al. (23), we also find a loss of TAMs with a pro-tumorigenic transcriptional profile upon STING agonist treatment. Interestingly, we report that the lack of this population in STING agonist treated tumors is dependent on intrinsic IFN-I signaling in monocytes, which to our knowledge had not been shown previously. Although we did not formally confirm the loss of TAM markers at the protein level, our findings are consistent with a recent study showing that intrinsic IFNAR1 signaling prevents the acquisition of immune suppressive features by myeloid-derived suppressor cells (43).

Based on CD64 and FcεR1 (MAR-1) staining, the inflammatory state of monocytic lineage cells in the LN and tumors was transient, peaking at day 1-2 and returning to baseline by day 5 post-ADU-S100 treatment. Monocytic lineage cells might re-acquire a pro-tumorigenic profile in absence of sustained IFN-I signaling. In a breast cancer model, depletion of TAMs with liposome clodronate during the first week of treatment with CAR-T cells and the STING agonist DMXAA reduced tumor control (23). However, depleting TAMs at later time points helped sustain tumor regression, indicating that TAMs reacquired a suppressive profile over time (23). In this case, an alternative approach to cell depletion could be the use of fractionated treatment regimens where STING agonists are given every 2-3 days, as this might sustain inflammatory states and prevent the resurgence of pro-tumorigenic TAMs.

Although cDC1 are specialized in the cross-presentation of exogenous antigens, other APC populations including monocytes, have been reported to acquire the ability to cross-prime CD8**^+^** T cells if appropriately stimulated (8, 44–47). However, the idea that infMCs can migrate to the LN and effectively cross-prime CD8**^+^** T cells has recently been challenged (29, 31). The characterization of a population of bona-fide cDC2s that express CD64 and stain positive for MAR-1 (infDC2s) suggested that the observed migration and priming ability of infMCs might have been attributable to contaminating cDCs. To accurately delineate infMCs from infDC2s in mice, it is now recommended to use CD26 and CD88 (29). Here, we report that CD26^-^CD88**^+^** Ly6C^hi^ MCs effectively take up tumor proteins and carry them to the LNs in response to intra-tumoral STING agonist administration. These cells were phenotypically and transcriptionally distinct from CD26**^+^**CD88^-^ infDCs, and expressed conventional monocyte genes such as *Csf1r*, *Ccr2*, *Itgam* and *Plac8*. A possible explanation for these conflicting findings is that we did not pre-gate on MHCII^hi^ cells in our analyses of infMCs. As infMCs in the LN express significantly lower levels of MHCII than cDCs, pre-gating on MHCII^hi^ cells might exclude them from the analysis.

Recently, CCR5 has been implicated in trafficking of monocytes to LNs (32). Consistently, we observed CCR5 protein expression on the monocytes. Moreover, *Ccr5* was identified as a monocyte but not DC expressed gene in the CITE-seq data, and the CCR5 antagonist maraviroc substantially reduced the trafficking of monocytes from the tumor to the dLN. We also provided evidence that MC in the LN substantially increase IFN-γ production by CD8 T cells through provision of IL-18, suggesting that the trafficking of monocytes to the LN is important in local T cell responses. Although not formally tested here, the presumption is that these cells then migrate back to the tumor to carry out their effector function. Indeed, it has been suggested that the dLN provides an important reservoir of differentiated T cells, protected from the inhibitory environment of the tumor, thus the delivery to the LN of tumor and IL-18 by monocytes may be relevant to the activation of T cells and tumor control (48).

Our data also showed that STING agonists induce immunological control of tumors at least in part through increased IL-18 production by monocytes. *Il18* was expressed by infMCs which have responded to IFN-I in the tumor and LN after STING agonist therapy. This is in line with previous studies demonstrating that IL-18 is transcriptionally regulated by STING and/or IFN-I signaling (49, 50). We confirmed the biological relevance of IL-18 induction in monocytes, by showing that in vivo neutralization of IL-18 reduced the efficacy of STING agonists in driving tumor control in *Ccr2^+/+^* but not *Ccr2^-/-^* mice. Of note, IFN-I dependent production of IL-18 by infMCs is also important for immune responses to viral, parasitic, or bacterial infections (6, 7). While *Il18* and intracellular IL-18 levels are upregulated in MCs upon exposure to IFN-Is, IL-18 requires proteolytic cleavage downstream of inflammasome activation to be secreted. There is evidence that the cGAS-STING-IFN-I axis activates the inflammasome in human and mouse MCs (51–53).

We also observed strong IFNAR1-dependent increases in MHC I and genes associated with Ag presentation in the CITE-seq data and we observed increased MHC I protein expression on the infMC in response to STING agonist therapy, consistent with increased peptide loading. However, *Ccr2^-/-^*mice which lack infMC, did not show impairment in the initial proliferation of adoptively transferred OT-I T cells compared to WT mice, arguing that infMCs are not significantly contributing to T cell priming in this model. These findings are consistent with the observation Batf3^+^ DCs are important for T cell priming during STING agonist therapy of cancer (54). In viral infection models, signals from monocyte-derived cells promote the post priming accumulation of T cells, with no detectable impact on initial cell proliferation over the first 5 days (4, 55). It is possible that low level of Ag presentation by the monocytes, while insufficient for priming CD8 T cells, could contribute to the interaction of the IL-18-producing monocytes with primed T cells in the LN, to augment effector function.

In conclusion, we have shown that intra-tumoral STING agonist therapy mediates at least part of its effects through infMCs. Mechanistically, STING agonists increased IFN-I signaling in monocytes, driving a unique inflammatory state that includes production of IL-18, which was important for CD8 T cell production of IFN-γ in the dLN, and for tumor control. Taken together, these findings highlight the importance of infMCs in STING agonist therapy of cancer. Monitoring STING agonist therapy in humans for induction of infMCs and IL-18 could help optimize such therapies.

## Acknowledgements

The authors thank Birinder Ghumman for technical assistance; Nathalie Simard and Janine Charron for assistance with fluorescence-activated cell sorting; Dr Slava Epelman and Rysa Zaman for breeding *Ifnar1^fl/fl^ Lyz2-Cre* mice; Dr Jean Gariépy for sharing the MC-38 cell line, and Drs. Sonya MacParland and Arthur Mortha for critical reading of the manuscript.

