## Supplemental figures 1-4 with legends for "STING agonists drive recruitment and intrinsic type I interferon responses in monocytic lineage cells for optimal anti-tumor immunity" (from another version of this preprint: version 1, 2024-12-11)

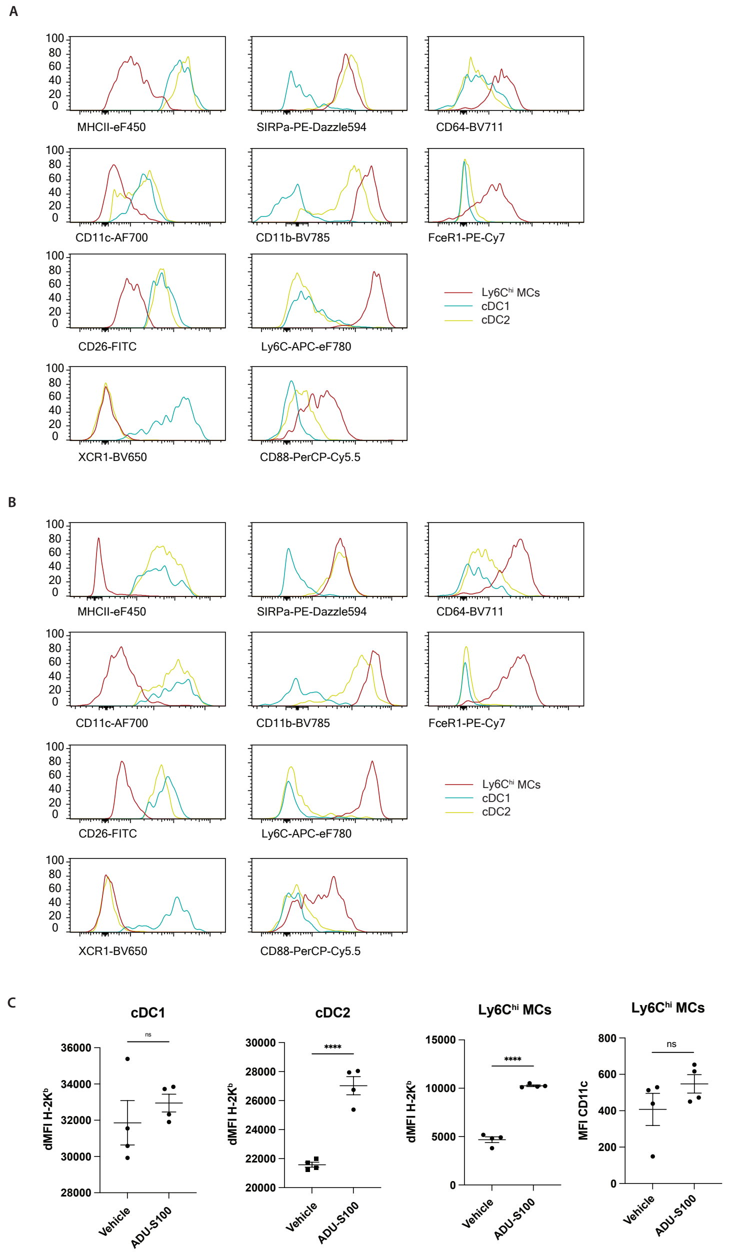


**Figure S1, supporting data for Figure 1. Cell surface marker expression on LN and tumor monocytes and DCs. (A-B)** Histograms depicting expression of indicated cell surface markers on Ly6C$^{hi}$ MCs, cDC1 and cDC2 in the lymph node 1-day post-treatment with ADU-S100 (A) or in the tumor (B) 2 days post-treatment with ADU-S100. Due to the low numbers of cDCs in the tumor day 2 post-treatment, histograms for cDC1 and cDC2 in (B) are from vehicle-treated mice. **(C)** Expression of MHC class I and CD11c in the TDLN 1-day post-treatment with vehicle or ADU-S100. Unpaired t-test (C) was performed.

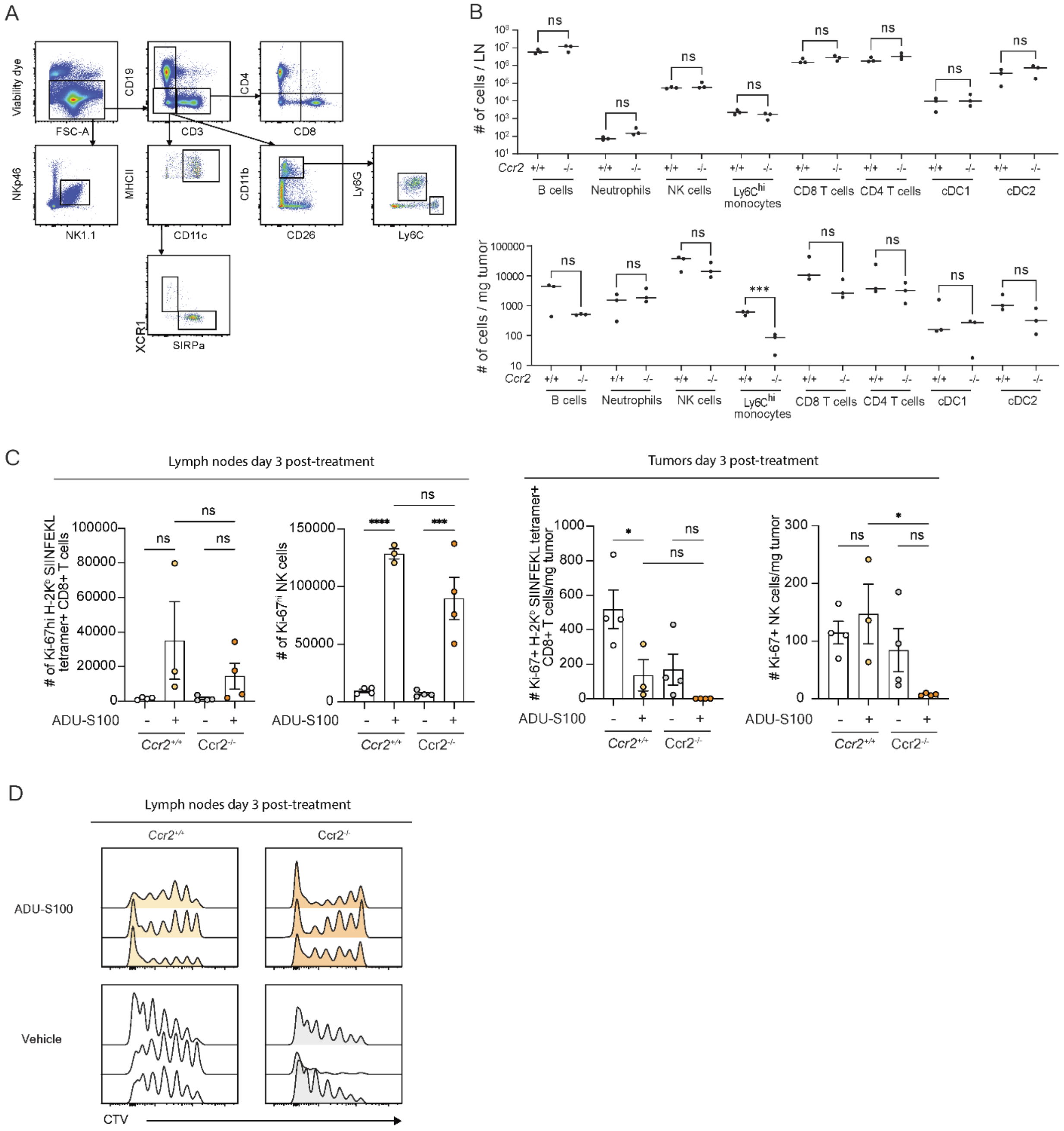

**Fig. S2. Supporting data for Figure 2.** (**A**) Gating strategy detecting cellular composition in *Ccr2*$^{+/+}$ and *Ccr2*$^{-/-}$ mice. (**B**). Comparison of cellular abundance *Ccr2*$^{+/+}$ and *Ccr2*$^{-/-}$ mice. (**C**). Absolute number of Ki-67+ tumor-specific T cells and NK cells in the TDLNs and tumor. (**D**). Histogram plots of adoptively transferred CellTrace$^{TM}$ violet (CTV) labelled OT-I cells in *Ccr2*$^{+/+}$ and *Ccr2*$^{-/-}$ mice on day 3 post-treatment in TDLN. Results shown are for 3 individual mice per condition from a single experiment and are representative of 2 similar experiments. Unpaired t-test (B), one-way ANOVA with Sidak's multiple comparisons test (C) was performed.

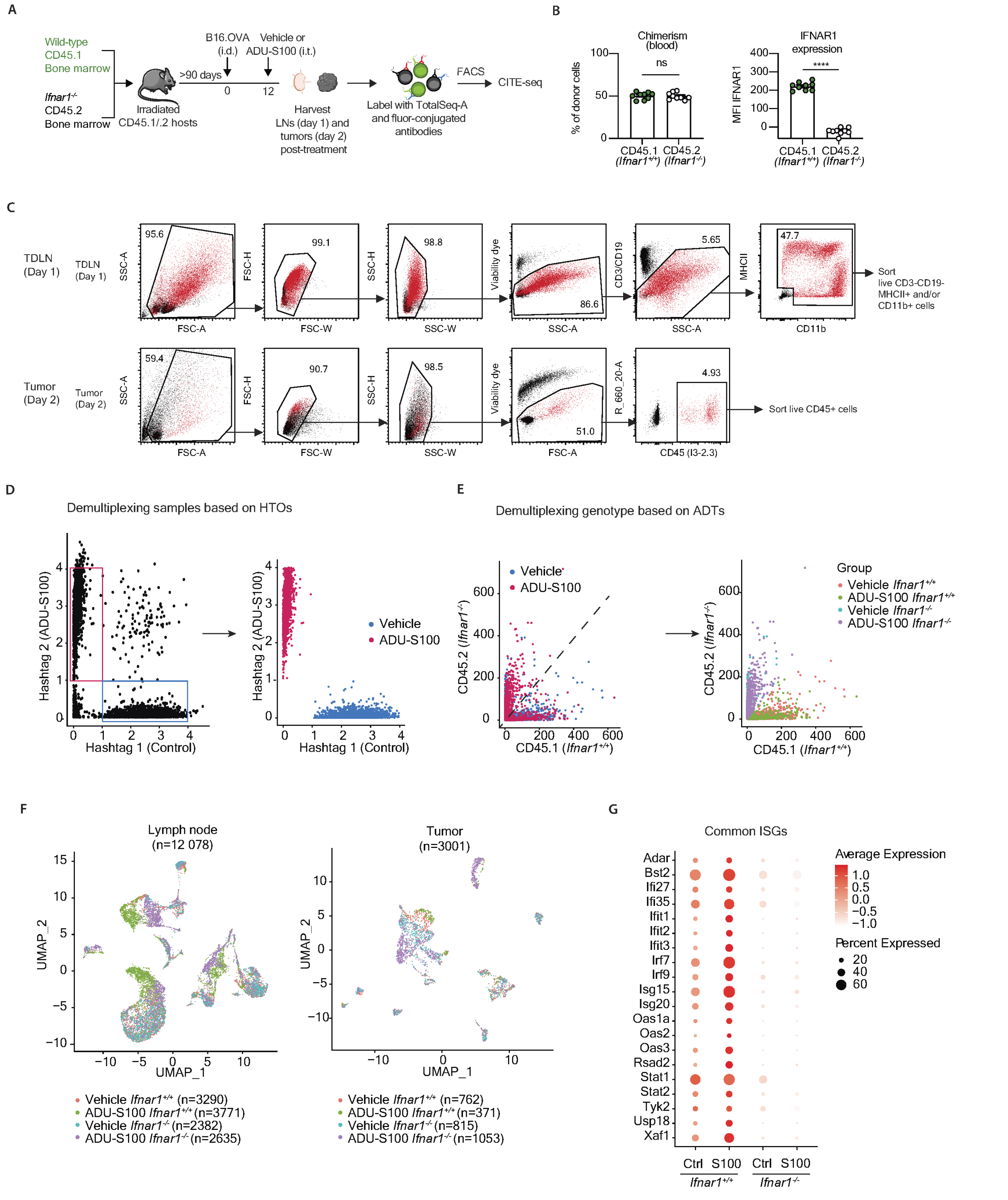

**Fig. S3**. **Supporting data for the CITE-sequencing experiment in Fig. 4. (A)** Experimental schematic. **(B)** Chimerism in blood CD45$^{+}$ cells. **(C)** Gating strategy for sorting of live CD3$^{-}$CD19$^{-}$ MHCII$^{+}$ and/or CD11b$^{+}$ cells from the LN and of CD45$^{+}$ cells from the tumors. **(D-E)**

Demultiplexing of samples based on hashtag oligos (HTOs) (**D**) or antibody-derived tags (ADTs)(E). **(F)** UMAP plots depicting total cells that passed quality control in the LN (left panel) and tumor (right panel) colored by treatment group x genotype. **(G)** Expression of common ISGs in total cells sequenced from the tumor validating the demultiplexing strategy.

A

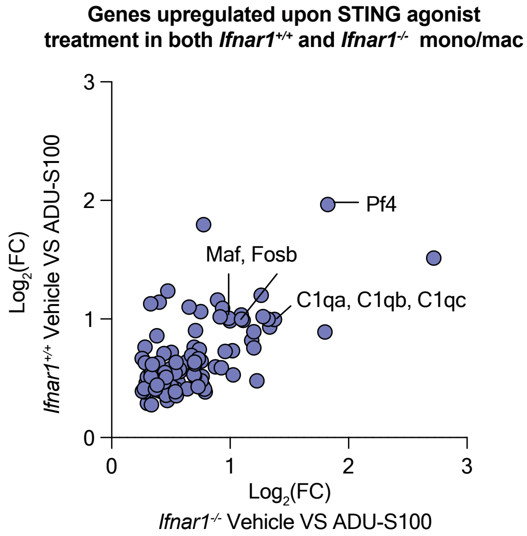


**Fig. S4. Effects of STING agonists on monocytes and macrophages that are independent of intrinsic IFNAR1 signaling. (A)** Genes significantly upregulated in monocytes and macrophages from ADU-S100 treated tumors compared to vehicle-treated tumors within the *Ifnar1*$^{+/+}$ and *Ifnar1*$^{-/-}$ compartment of mixed bone marrow chimeras. Shown are the genes that are significantly upregulated in both the *Ifnar1*$^{+/+}$ and *Ifnar1*$^{-/-}$ compartments following STING agonist therapy, suggesting that they are regulated by STING agonists independently of intrinsic IFNAR signaling.
